# A nearly optimal sequential testing approach to permutation-based association testing

**DOI:** 10.1101/186361

**Authors:** Julian Hecker, Ingo Ruczinski, Brent Coull, Christoph Lange

## Abstract

The following technical report describes the technical details for the implementation of a sequential testing approach to permutation-based association testing in whole-genome sequencing studies. The sequential testing approach enables to control the probability of a type 1 and type 2 error at arbitrary small pre-specified levels and approaches the theoretical minimum of expected number of required permutations as these levels go to zero.

---

In practice, since it is not feasible to go through all permutations of a genetic data set, the permutation-based p-value is usually estimated from a large number of random permutations.

The procedure of the re-calculation of the association test statistic for permuted data and comparison with the observed test statistic can be described by a sequence *x*_1_, *x*_2_,…, where *x*_*i*_ = 1 if the *i*-th permuted test statistic is larger or equal than the observed statistic and *x*_*i*_ = 0 otherwise. Denote the true and unknown association p-value, computed by the evaluation of all permutations (not feasible in practice), by *θ*. The scientific interest is mainly summarized by the question if θ ≤ *p*_1_, for a pre-specified significance level *p*_1_. For example, *p*_1_= 5 · 10^−8^ in classical GWAS. Given the extremely large number of possible permutations and assuming an appropriate generation of random permutations, we interprete the sequence *x*_1_, *x*_2_,… as independent, identically distributed Bernoulli random variables with success probability *θ*. In the following, we describe how significance testing can be performed efficiently by a sequential testing approach.

## Setting

Let (Ω, *Ƒ* P_*θ*_) be a probability space *x*_1_, *x*_2_, … ∽ Bernoulli(*θ*), a sequence of independent identically distributed Bernoulli random variables with success parameter *θ*, and *Ƒ*_*n*_ = σ (*x*_1_, …, *x*_*n*_) ⊂ *Ƒ, for n* = 1,2,…. 0 ≤ *θ* ≤ 1 is the true p-value, that can computed (theoretically) by evaluating all permutations of the data set. We extended the standard Bernoulli distribution for *θ* ∈ (0,1) to the inclusion of the extreme cases *θ* =1 and *θ* = 0.

We would like to differentiate between two hypotheses:

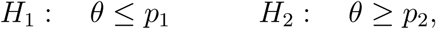

where *p*_2_ − *p*_1_ = *d* > 0, *p*_1_ > 0 and *p*_2_ < 0.5. In practice, we choose, for example, *p*_1_ = 5 · 10^−8^ (genome-wide significane in classical GWAS), *d* = 10^−8^(resolution level of 10^8^ permutations). *d* is chosen small and affects the worst case expected run time, as described below. The interval (*p*_1_, *p*_2_) is the so-called indifference zone, where both hypotheses are plausible.

## Sequential testing framework: Corresponding objects and results

We utilize the work of Pavlov (1991) [1] and Tartakovsky (2014) [2] for sequential testing theory. The following strategies and results are strongly related to the work in [1] and adapt these results from the general setting to our specific scenario. In particular, we show that our estimator for *θ* is an appropriate choice and deal with the problem of the degenerated cases *θ* = 1 or *θ* = 0.

Let *D*_1_ ≔ [0,*p*_1_] and *D*_2_ ≔[*p*_2_, 1]. Let (*α*_1_, *α*_2_) = (*αt*_1_, *αt*_2_) for positive *t*_1_, *t*_2_, *α* such that *α*_1_ + *α*_2_ < 1.

Introduce

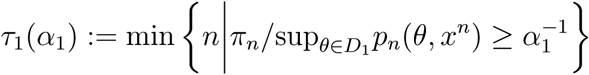

and

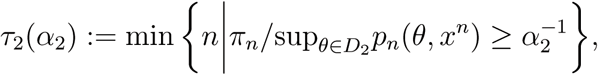

where 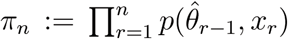 with π_1_ ≔ 0.5 and *p*_*n*_(*θ, x*^*n*^) = 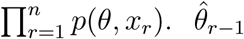 is an estimate of *θ* which depends only on the first *r* − 1 observations and is specified by 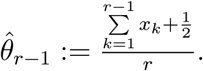

A decision test is described by *Ƒ*_*n*_-stopping time *N* and a *Ƒ*_*N*_-measurable function *δ*, which can take the values 1 and 2.

We define the decision test for our approach as

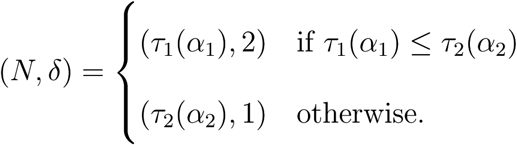

Denote by *ρ*(θ_1_, θ_2_) the Kullback-Leibler Distance for the Bernoulli distribution, defined by 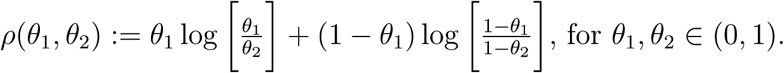

From our choice for the decision test, we get the following results.

### Theorem

*1.) For the error probabilities, we have*

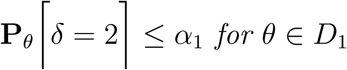

*and*

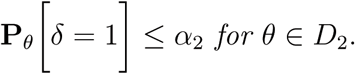

*2.) For the expected number of permutations, we have*

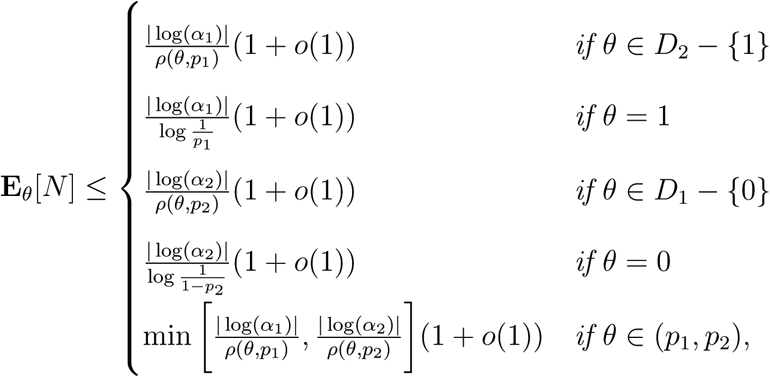

as α → 0.

*3.) Let K*(*α,t*_1_, *t*_2_) *be the class of all decision tests* (*N*′, *δ*′) *such that* 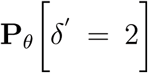 ≤ *αt*_1_ *for θ ∈ D*_1_ *and* 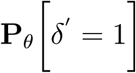 ≤ *αt*_2_ for θ ∈ *D*_2_, *then*

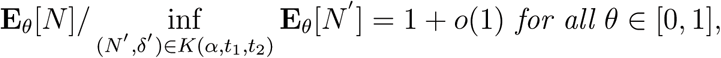

as *α* → 0.

### Remark 1

*Note that, for given p*_2_ ∈ (0,1), *we have* 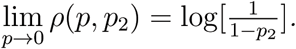 *This shows that the results for θ = 0 and θ=1 are the natural extensions*.

## Proof of the Theorem

The proof of the Theorem is strongly related to the derivations in [1]. One important difference is that Pavlov derived uniform bounds, whereas our estimates will depend on *θ*. We use the explicit form of 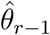 and show how we can deal with the cases *θ*=1 and *θ* = 0.

### Lemma 1

*For all* θ ∈ *D*_1_

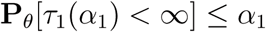

*and for all θ ∈ D*_2_

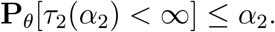

*Proof*. We use the same argumentation as in [1]. In our setting, for any *θ ∈* (0,1), the process 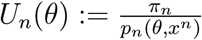 forms a non-negative martingale with respect to *Ƒ*_*n*_. In addition, we have *E*_θ_[*U*_*n*_(θ)] = 1, since π_1_ ≔ ½. As in [1], introduce *v*_*θ*_ ≔ min 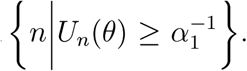 By Doobs inequality, we have 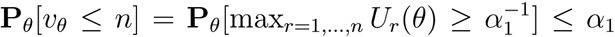 for all *n* and so P_*θ*_[*v*_*θ*_ < ∞] ≤ α_1_. For *θ ∈ D*_1_ with *θ* ≠ 0, we haveτ_1_(α_1_) ≥ *v*_θ_. Therefore, we get

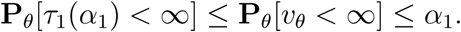

For *θ* = 0, obviously P_θ_[τ_1_(α_1_) < ∞] = 0. Same argumentation for *θ* ∈ *D*_2_ and P_*θ*_[τ_2_(α_2_) < ∞].□

We define 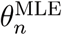 as the ordinary maximum likelihood estimator for 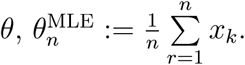

Furthermore, 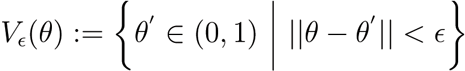 and 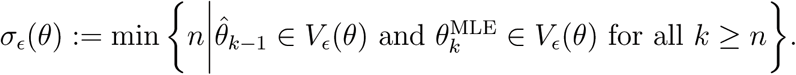

### Lemma 2

*For all θ ∈* (0,1) *and* ε > 0 *such that V*_ε_(θ) ⊂ (0,1),

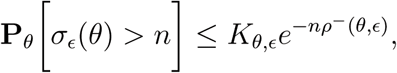

*for all n, where ρ*^-^(*θ,ε*) ≔ min[*ρθ + ε, θ*), *ρθ + ε, θ*)].

*Proof*. Since we have an explicit form for the estimators 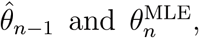 the proof is straightforward. Fix *θ* ∈ (0,1) and ε > 0 such that *V*_ε_(θ) ⊂ (0,1). Define 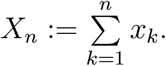

Start with

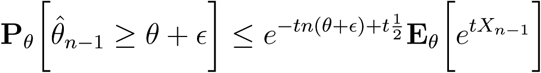

for all *t* > 0.

From the classical proof of the Chernoff-Hoeffding bound using moment-generating functions, we know that there is a *t*\* > 0, which depends on *θ* and ε, such that

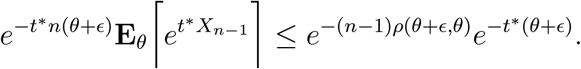

In addition,

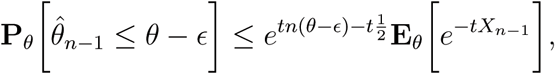

for *t >* 0. Analogous argumentation shows the estimate for both estimators. The statement of the Lemma follows from

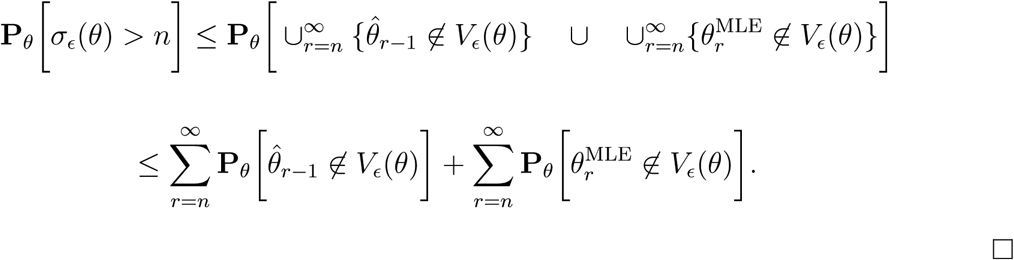

Define 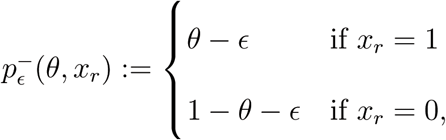 for an appropriate ε > 0, such that *θ* - ε > 0 and 1 − *θ* − ε > 0.

### Lemma 3

*Let θ ∈* (0,1), θ_0_ ∈ (0,1) *and δ >* 0. *If ε > 0 is chosen small enough such that*

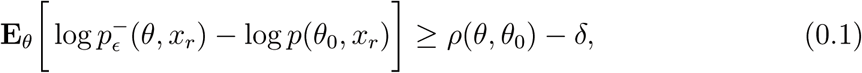

*then*

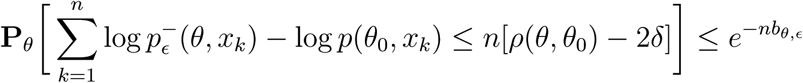

*for all n, where b*_θ,ε_ > 0.

*Proof*. The estimate e^−nb_θ,ε_^ depends on *θ* and ε, in opposite to the estimate in [1]. We proceed as in [1], but we use the explicit form of the Bernoulli distribution. Let *t >* 0, define

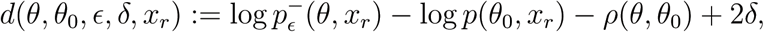

and easily compute

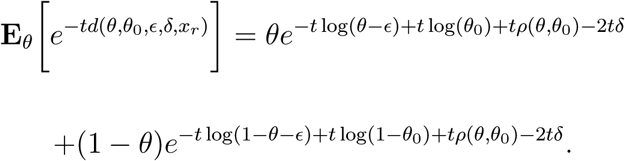

From here we can use the argumentation as in the proof of Lemma 5.1 in [1]. □

### Lemma 4

*Let* ε > 0 *and define*

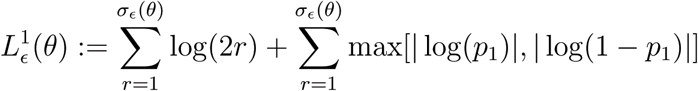

*and*

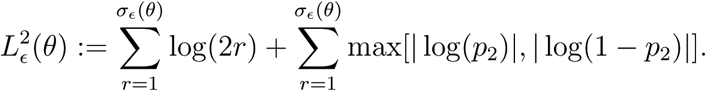

*Then*,

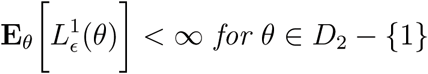

*and*

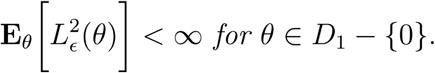

*Proof*. For *θ ∈ D*_2_ − {1}, we estimate

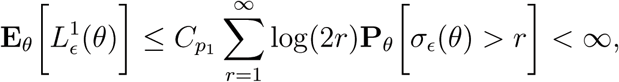

by the ratio test for infinite series, Lemma 2 and the properties of the geometric sum. Same argumentation for 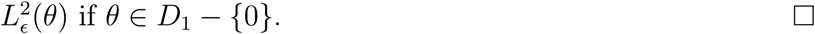

*Proof of the Theorem*. We start with 2.)

Let *θ ∈ D*_2_ − {1} and 0 < *δ* < ρ(*θ,p*_1_). Choose ε > 0 such that *V*_ε_(θ) ∈ (*p*_2_, 1) and such that

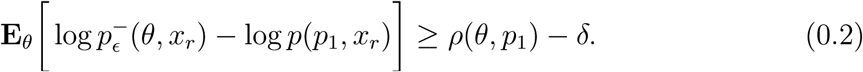

Then, we can estimate

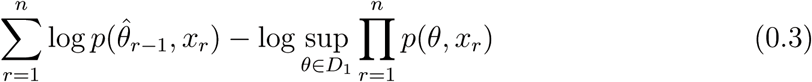

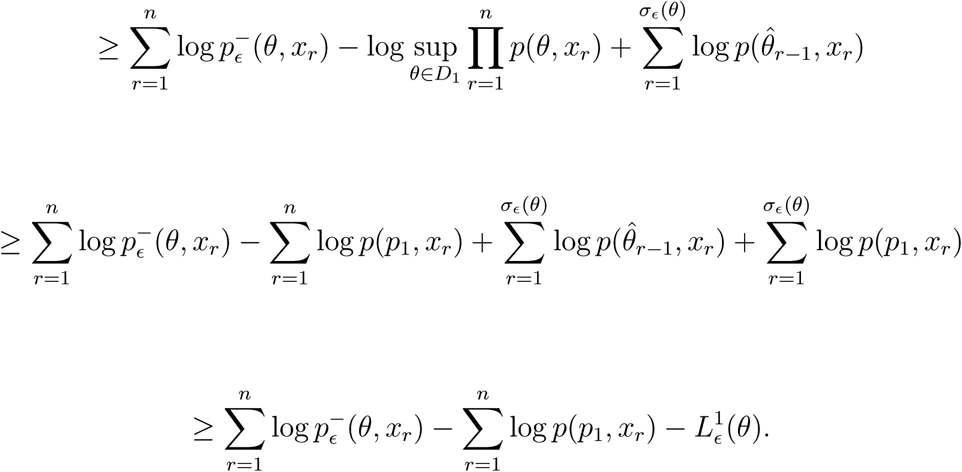

Define

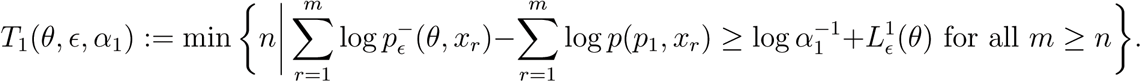

From this point, with the same argumentation as in the proof of Lemma 5.6 in [1], using Lemma 3, it follows

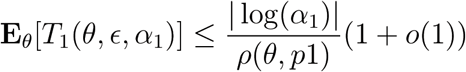

as *α* → 0. The same can be shown for the analogously defined *T*_2_(*θ,ε, α*_2_). This together with Eq.(0.3) concludes the claim regarding the expected number of permutations for *θ ∈ D*_1_ − {0} and *θ ∈ D*_2_ − {1}. For *θ* ∈ (*p*_1_, *p*_2_) the claim follows since the ratio between α_1_ and α_2_ is assumed to be fixed and equal to 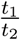 as α → 0.

In the scenario *θ* = 1, we have a deterministic setting with *x*_*r*_ = 1 for all *r*. Then, *N* is determined by

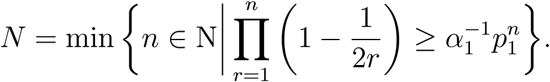

Furthermore,

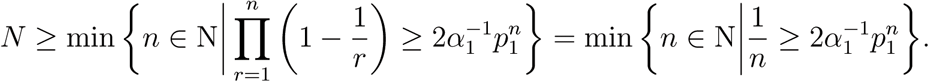

If we analyze min 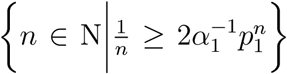 using the Lambert W function and the corresponding asymptotic expansions, we can conclude

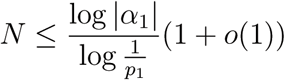

as α → 0. Exactly the same argumentation shows the desired statement for *θ* = 0.

1.) We showed for all *θ ∈* [0,1] that 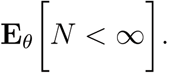 This implies 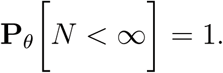 Therefore, for *θ ∈ D*_1_,

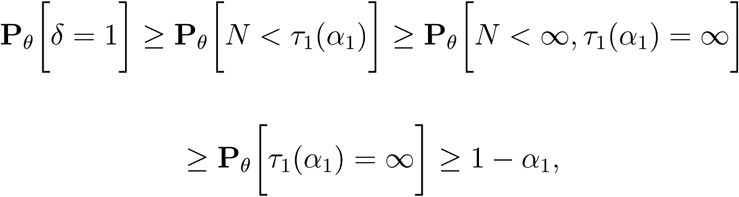

leading to

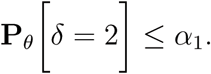

Analogously, 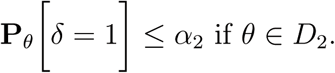

3.) From Lemma 3.2 in [2], we obtain exactly the same bounds as stated in 2.) for θ ∈ (0, 1) as lower bounds for N′ if (*N*′, *δ*′) *∈ K (α, t*_1_, *t*_2_). Thus, only the cases *θ* = 1 and θ = 0 are missing. Consider *θ*=1 and assume there is a decision test in *K*(*α,t*_1_, *t*_2_) such that 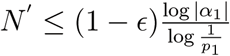 for any ε > 0. Then, we have

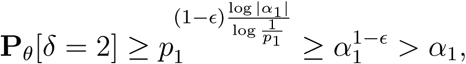

for *θ* = *p*_1_ ∈ *D*_1_, a contradiction. The same argument works in the case *9 = 0* and this concludes the proof of the statement 3.) □

